# Therapeutic base and prime editing of *COL7A1* mutations in recessive dystrophic epidermolysis bullosa

**DOI:** 10.1101/2021.07.12.452037

**Authors:** Sung-Ah Hong, Song-Ee Kim, A-young Lee, Gue-Ho Hwang, Jong Hoon Kim, Hiroaki Iwata, Soo-Chan Kim, Sangsu Bae, Sang Eun Lee

## Abstract

Recessive dystrophic epidermolysis bullosa (RDEB) is a severe skin fragility disorder caused by loss-of-function mutations in the *COL7A1* gene, which encodes type VII collagen (C7), a protein that functions in skin adherence. From 36 Korean RDEB patients, we identified a total of 69 pathogenic mutations (40 variants without recurrence), including point mutations (72.5%) and insertion/deletion mutations (27.5%). We used base and prime editing to correct mutations in fibroblasts from two patients (Pat1, who carried a c.3631C>T mutation in one allele, and Pat2, who carried a c.2005C>T mutation in one allele). We applied adenine base editors (ABEs) to correct the pathogenic mutation or to bypass a premature stop codon in Pat1-derived primary fibroblasts. To expand the targeting scope, we also utilized prime editors (PEs) to correct the mutations in Pat1- and Pat2-derived fibroblasts. Ultimately, we found that both ABE- and PE-mediated correction of *COL7A1* mutations restored full-length C7 expression, reversed the impaired adhesion and proliferation exhibited by the patient-derived fibroblasts, and, following transfer of edited patient-derived fibroblasts into the skin of immunodeficient mice, led to C7 deposition within the dermal-epidermal junction. These results suggest that base and prime editing could be feasible strategies for *ex vivo* gene editing to treat RDEB.

## INTRODUCTION

Epidermolysis bullosa (EB) is a heterogeneous group of genodermatoses characterized by mucocutaneous fragility. Recessive dystrophic EB (RDEB), one of the most severe subtypes of EB, results from biallelic mutations in the *COL7A1* gene, which encodes the alpha 1 chain of type VII collagen (C7). C7 is a key component of anchoring fibrils, which create a strong attachment between the epidermis and dermis. Loss of C7 causes extensive mucocutaneous blistering, scarring, and extracutaneous complications, leading to considerable morbidity and occasional mortality.^[1]^ Therefore, effective treatments are urgently needed. To date, various therapeutic strategies, including protein replacement,^[2]^ disease-modifying drugs,^[3]^ and allogeneic cell-based therapies using fibroblasts,^[4]^ mesenchymal stromal cells,^[5]^ and bone marrow transplantation^[6]^ have been studied, but a complete cure is not achievable with those current approaches. As a potential long-lasting therapeutic option, remarkable progress has been made in gene therapy that aimed to transfer the normal *COL7A1* gene into deficient cells from RDEB patients. In this strategy, it is advantageous to use autologous cells for gene transfer. Early phase clinical trials using viral vectors such as lentivirus and retrovirus^[7]^ to transduce *COL7A1* into autologous fibroblasts and keratinocytes have resulted in C7 restoration in the treated skin of some RDEB patients for more than 1 or 2 years after treatment. Despite these promising results, such viral vector-based gene therapy has potential concerns: i) random integration of viral vectors into the host genome, ii) continued expression of aberrant transcripts from the endogenous *COL7A1* gene, and iii) different levels of constitutive expression from the virus-delivered exogenous gene regardless of the cellular environment.

To overcome these limitations, therapeutic editing of the endogenous *COL7A1* gene in patients’ autologous cells via genome editing tools has been suggested for RDEB treatment.^[8]^ Conventional CRISPR nucleases rely on double-strand breaks (DSBs) in the target DNA, which are repaired by one of the cell’s repair systems, such as non-homologous end joining (NHEJ) or homology-directed repair (HDR).^[9]^ Previously, several groups demonstrated the feasibility of correcting the reading frame^[8e, 8f]^ or skipping a mutant exon^[8g-i]^ in mutant *COL7A1* using CRISPR-coupled NHEJ repair. However, these strategies have limited value for the correction of point mutations, the most common type of mutation in RDEB. In contrast, the use of HDR enables the precise correction of point mutations, but its low editing efficacy, requirement for donor templates, and limited activity in non-dividing cells are obstacles for HDR-mediated approaches. Furthermore, recent studies have revealed that CRISPR nuclease-mediated DSBs can induce unwanted large deletions, chromosomal rearrangements,^[10]^ and a p53-mediated DNA damage response^[11]^ that results in cell death, potentially inhibiting further clinical applications.

To bypass such risks, newly developed tools that generate few DSBs, such as base editors (BEs) and prime editors (PEs), can be used.^[12]^ BEs, which include cytosine base editors (CBEs)^[13]^ and adenine base editors (ABEs)^[14]^, can convert one target nucleotide into another, C-to-T or A-to-G, by catalyzing cytosine or adenine deamination, respectively. A recent report described an ABE-mediated strategy in which two nonsense mutations in *COL7A1* were corrected *ex vivo* in RDEB patient-derived fibroblasts and induced pluripotent stem cells, resulting in therapeutic effects such as C7 restoration.^[15]^ However, despite their therapeutic potential, BEs have limited ability to correct small insertion and deletion (indel) mutations or transversion mutations such as C-to-G/A and A-to-C/T. Alternatively, PEs can generate all types of substitutions and indels within about a 40-bp sequence. A practical version of PE, PE2, which consists of a Cas9 nickase (nCas9) that contains a H840A mutation and an engineered reverse transcriptase, is recruited to the target site by a prime editing guide RNA (pegRNA).^[16]^ The pegRNA is composed of a standard single-guide RNA (sgRNA) and an extension sequence at the 3’ end that includes a primer binding site (PBS) and a reverse transcription template (RTT) that encodes the desired correction. To maximize PE efficacy, PE3 employs an additional nicking sgRNA (ngRNA) for inducing a second nick in the non-edited strand. However, prime editing has not yet been demonstrated for RDEB treatment.

In this study, we established a *COL7A1* mutation database from a large cohort of South Korean patients with RDEB and analyzed the percentage of mutations that are potentially targetable by BEs and PEs. We then applied either ABE or PE3 to correct the mutations in primary fibroblasts from two patients with highly recurrent *COL7A1* mutations. We further transplanted the ABE-/PE-corrected primary fibroblasts into immunodeficient mice and observed strong linear deposition of human type VII collagen (C7) at the dermal-epidermal junction (DEJ), supporting the therapeutic potential of *ex vivo* ABE- or PE-mediated gene editing for treating RDEB.

## RESULTS

### Establishment and analysis of a *COL7A1* mutation database specific for Korean RDEB patients

Using an *in silico* approach, we first inspected all known RDEB-associated *COL7A1* variations among the world-wide population of patients, and determined which ones would be targetable with BEs and PEs. According to the *COL7A1* variants database (http://www.col7a1-database.info), a total of 810 *COL7A1* gene variants causing RDEB are currently registered, of which 23.6% are indel mutations and 76.4% are point mutations, including nonsense, missense, synonymous, and intron mutations (**Figure 1A**). Among the *COL7A1* point mutations, 2.9% (i.e., A>G or T>C) and 26.2% (i.e., G>A or C>T), respectively, can theoretically be corrected by CBEs and ABEs derived from SpCas9 (Cas9 from *Streptococcus pyogenes*), which recognize a canonical 5’-NGG-3’ protospacer adjacent motif (PAM). When NG-PAM-targetable BEs are used instead, 6.0% and 37.6% of mutations are covered by CBEs and ABEs, respectively.^[17]^ In contrast, given that PEs can correct all types of point mutations as well as indel mutations, 96.2% of the mutations can potentially be corrected by NG-PAM-targetable PEs (**Figure 1A**).

**Figure 1.**
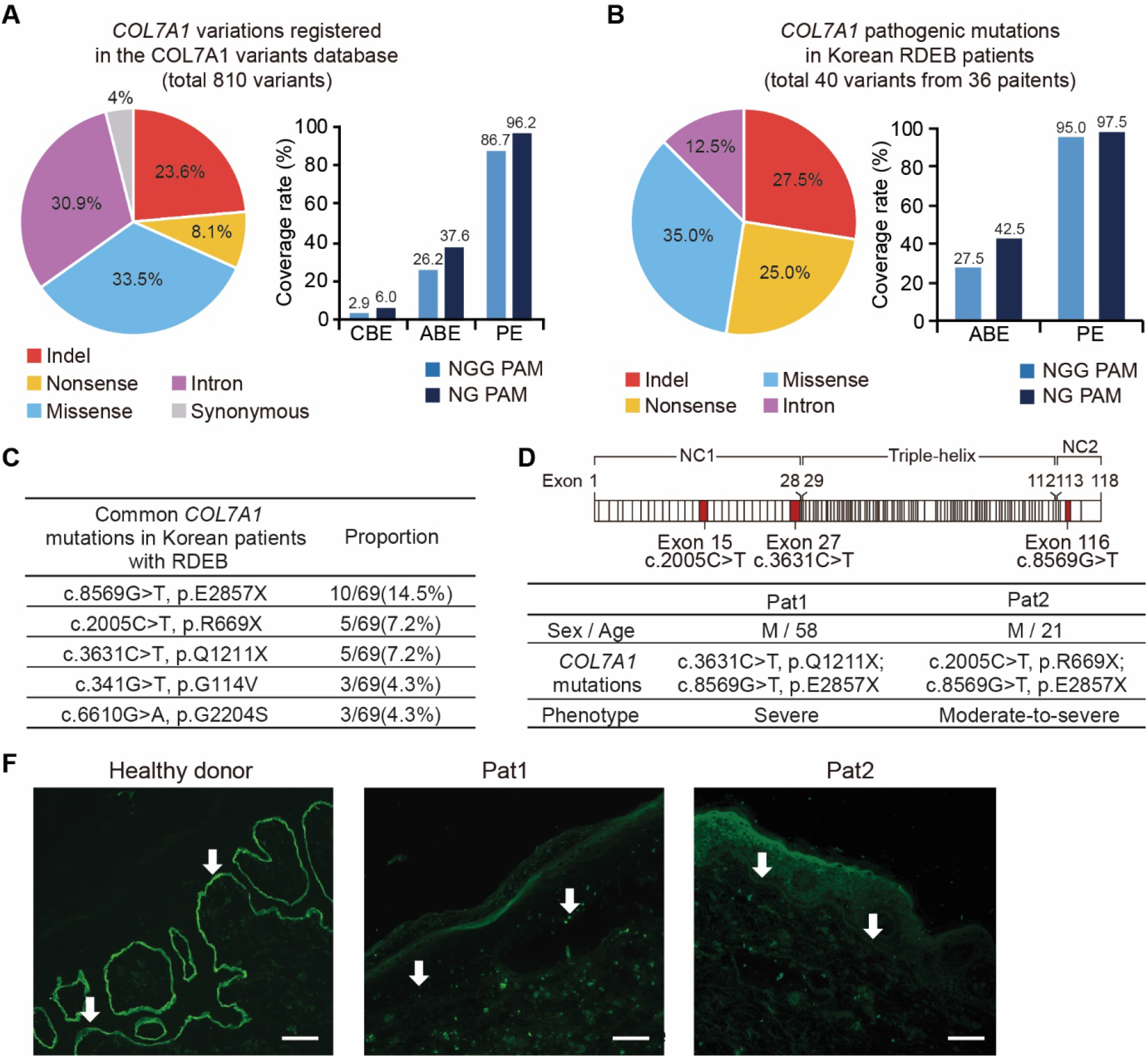
Establishment of a *COL7A1* mutation database specific for Korean RDEB patients and analysis of Korean and world-wide databases. (a) The full spectrum of pathogenic *COL7A1* mutations reported to date in DEB patients. Frequencies (%) of *COL7A1* alleles that can theoretically be corrected with CBE, ABE, and PE constructed with SpCas9 (NGG-PAM) or its variant with a relaxed PAM requirement (NG-PAM). (b) Mutational analysis of *COL7A1* in a large cohort of Korean RDEB patients. Frequencies (%) of *COL7A1* alleles that can theoretically be corrected with ABE and PE recognizing NGG- or NG-PAMs. (c) The five most frequent pathogenic *COL7A1* mutations in our database among the 36 Korean RDEB patients with 69 pathogenic mutations. (d) Information about the two RDEB patients enrolled in this study and a schematic representation of procollagen VII showing the locations of the *COL7A1* mutations identified in these patients. (e) Immunofluorescence to visualize C7 was performed on skin samples from Pat1, Pat2, and healthy controls using polyclonal rabbit anti-COL7 antibody. White arrows point at the DEJ. Representative images are shown. Scale bars represent 50 μm.

Similarly, we further investigated the editing scope of both BEs and PEs for correcting pathogenic *COL7A1* mutations found in Korean patients suffering from RDEB. To this end, using information from the only EB referral center in South Korea, we established the largest database of *COL7A1* mutations in Korean RDEB patients and identified a total of 69 pathogenic mutations (40 variants without recurrence) from a total of 36 patients. Of the 40 mutations, 72.5% were point mutations, including missense (35.0%), nonsense (25.0%), and intron (12.5%) mutations, whereas 27.5% were indel mutations (**Figure 1B** and **Table S1**). Among the point mutations, 27.5% can theoretically be corrected by NGG-PAM-targetable ABEs, and 42.5% can theoretically be corrected by NG-PAM-targetable ABEs. When PEs were considered, we found that 97.5% of the mutations would be covered by NG-PAM-targetable PEs (**Figure 1B**), consistent with the situation for the world-wide population of patients.

In the Korean RDEB database, c.8569G>T (p.E2857X), c.2005C>T (p.R669X), and c.3631C>T (p.Q1211X) were the most recurrent RDEB-causing *COL7A1* mutations, representing 14.5% (10/69), 7.2% (5/69), and 7.2% (5/69) of the mutant alleles, respectively (**Figure 1C**). Among these, c.2005C>T in exon 15 and c.3631C>T in exon 27 affect the amino-terminal non-collagenous NC-1 domain and have been reported to induce nonsense‐mediated decay of COL7A1 transcripts.^[18]^ In addition, these two nonsense mutations have been reported to be responsible for severe generalized RDEB (**Figure 1D**).^[18b,19]^ Therefore, we focused on these two nonsense mutations in our cohort as targets for correction via ABEs or PEs. Two patients with moderate-to-severe RDEB who had compound heterozygous *COL7A1* nonsense mutations were enrolled in this study; patient #1 (Pat1, hereafter) carried c.3631C>T and c.8569G>T mutations and patient #2 (Pat2, hereafter) carried c.2005C>T and c.8569G>T mutations (**Figure 1D**). Skin biopsies from both patients showed only trace staining of C7 by immunofluorescence microscopy using antibodies against the NC-1 domain, whereas skin samples from healthy donors showed clear C7 staining at the DEJ (**Figure 1F**).

### Adenine base editing for *COL7A1* gene correction in Pat1-derived fibroblasts

To correct the C>T nonsense mutations, we first used the ABE system and prepared an optimized version of ABE7.10, named ABEmax.^[20]^ We used two strategies to rescue *COL7A1* gene function in Pat1-derived fibroblasts: i) direct correction of the mutated nucleotide (i.e., c.3631C>T) using sgRNA#1 (Pat1-sg1, hereafter) and ii) readthrough of the premature stop codon (PTC) using a method called CRISPR-pass,^[21]^ which involved editing the neighboring sequences using sgRNA#2 (Pat1-sg2, hereafter) (**Figure 2A**). Using Pat1-sg1, correction of the pathogenic mutation at position 5 (counting from the 5’ end of the target sequence) in the protospacer sequence would occur by conversion of adenine to guanine in the template strand, resulting in T-to-C correction on the coding strand. Using Pat1-sg2, the PTC (5’-TAG-3’) caused by c.3631C>T can be converted to 5’-TGG-3’, which will be translated to tryptophan (Trp), leading to restoration of the *COL7A1* reading frame. Because this amino acid change was predicted to have no deleterious effects on C7 (PROVEAN score < -2.5; PredictProtein score >50), we hypothesized that Pat1-sg2-induced PTC readthrough could contribute to C7 restoration despite this amino acid change.

**Figure 2.**
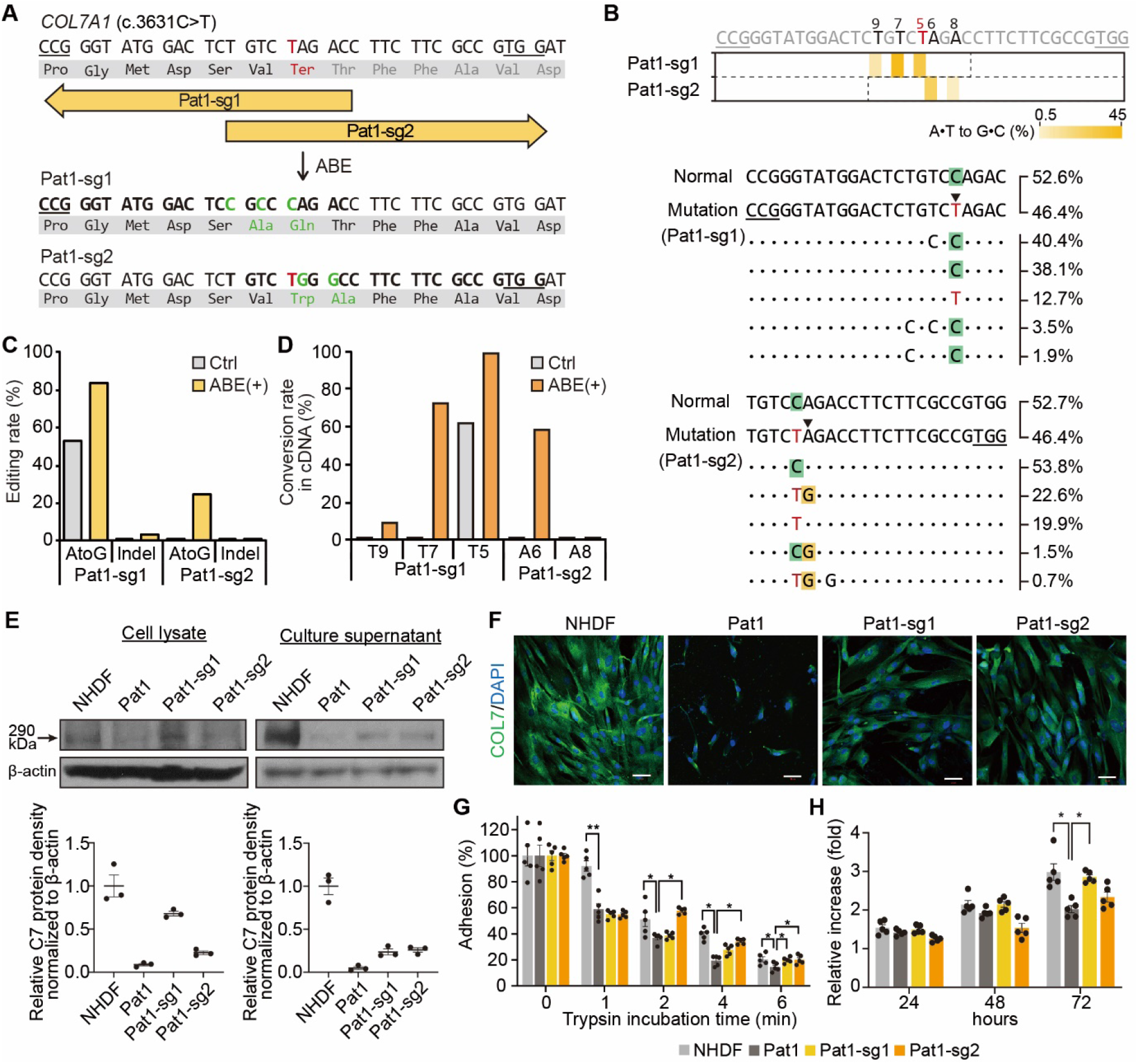
Correction of c.3631C>T (Q1211X) in *COL7A1* using ABEs in primary RDEB fibroblasts. (a) Schematic diagram of the ABE target sequence in exon 27 of the *COL7A1* gene containing a C>T nonsense mutation (c.3631C>T, p.Q1211X). The mutant sequence is shown in red. Target sequences are highlighted in bold type, and PAM sequences are underlined. A•T to G•C conversions in the ABE editing window are shown in green. (b) Heatmap visualizing A•T to G•C conversion rates analyzed by high-throughput sequencing (top) and the sequences at the target sites together with their proportions (middle and bottom). The five most common sequences in ABE-treated RDEB fibroblasts are shown, and the frequencies of normal and mutated allelesin the non-edited patient-derived fibroblasts are shown at the top of the panel. PAM sequences are underlined, and the target A•T is indicated by an arrowhead. (c, d). Conversion rates calculated by deep sequencing of genomic DNA (c) and mRNA (d) from RDEB fibroblasts treated with different sgRNAs. (e) Western blots to measure C7 abundance in cell lysates and culture supernatants using β-actin as an internal control. Protein band densities from three independent experiments are presented as bar graphs. Each density value was normalized to the β-actin value and expressed relative to the value in NHDFs. Data are mean ± SEM. (f) Immunofluorescence staining to visualize the C7 protein (green) in NHDFs, non-edited RDEB fibroblasts from Pat1, and ABE-treated RDEB fibroblasts. Nuclei were stained with DAPI (blue). Scale bars, 50 μm. (g) Trypsin-based cell detachment assay. Cell adhesion is represented as the percentage of cells that remain attached after the indicated period of trypsin treatment. Five independent experiments were performed. Data are mean ± SEM. *P < 0.05, ^**^P < 0.01. (h) The proliferation of NHDFs, non-edited fibroblasts from Pat1, and ABE-treated RDEB fibroblasts was evaluated by the WST-1 assay. The ratio of the absorbance at 24, 48, and 72 hours to that at 0 hours is shown. Five independent experiments were performed. Data are mean ± SEM. ^*^P < 0.05, ^**^P < 0.01.

Pat1-derived fibroblasts were then transfected with the ABEmax-encoding plasmid and each sgRNA-encoding plasmid by electroporation and harvested after 3 to 7 days. Genomic DNA was isolated from the bulk population of cells and subjected to high-throughput sequencing for the assessment of base editing outcomes. The sequencing results showed that, through strategy (i) using Pat1-sg1, the target T (T5) was efficiently converted to C at a frequency of 30.6%, whereas bystander Ts (T7 and T9) were also edited at frequencies of 44.5% and 5.5%, respectively (**Figure 2B** and **2C**). On the other hand, through strategy (ii) using Pat1-sg2, the target A (A6) and a bystander A (A8) were converted at frequencies of 24.6% and 1.8%, respectively (**Figure 2B** and **2C**). The frequencies of indels generated by ABE/Pat1-sg1 and ABE/Pat1-sg2 were 3.3% and 0.1%, respectively (**Figure 2C**). We further assessed the frequencies of *COL7A1* editing outcomes at the mRNA level using complementary DNAs (cDNAs). We found that the target sequences were edited at rates that were higher than that in genomic DNA, similar to findings from previous studies (**Figure 2D** and **S1A**).^[15, 22]^

Next, we evaluated C7 expression in ABE-treated RDEB fibroblasts from Pat1. Western blot analysis of bulk populations of such cells revealed the restoration of the full length C7 protein, at levels that were up to 68% (Pat1-sg1) and 23% (Pat1-sg2) of the the level in normal human dermal fibroblasts (NHDFs), whereas uncorrected cells showed barely detectable C7 protein (**Figure 2E**). The amount of C7 released into the culture supernatant of the RDEB fibroblasts was also increased following ABE treatment, to up to 19% (Pat1-sg1) and 22% (Pat1-sg2) of the levels seen in the NHDF supernatant (**Figure 2E**). Immunocytochemistry of C7 confirmed these findings and revealed increased C7 protein expression in the cytoplasm of ABE-treated RDEB fibroblasts (**Figure 2F**). It was previously reported that RDEB fibroblasts exhibited decreased adhesion ability due to C7 deficiency, and that viral vector-mediated transduction of the full-length human *COL7A1* gene restored their adhesion capacity.^[23]^ Thus, we further evaluated the adhesion properties of the ABE-treated RDEB fibroblasts using a trypsin-based cell detachment assay. Whereas uncorrected RDEB fibroblasts showed poor cell adhesion, with 59%, 37%, and 19% of cells adhering at 1, 2, and 4 minutes after trypsin treatment compared to 92%, 51%, and 40% of NHDFs, ABE-treated RDEB fibroblasts (Pat1-sg1) showed a 21%, 15%, and 6% increase in cell adhesion at 2, 4, and 6 minutes compared to untreated RDEB fibroblasts (**Figure 2G**). We also tested the effect of *COL7A1* correction on the ability of RDEB fibroblasts to proliferate using a mitochondrial activity assay (WST-1 assay). The RDEB fibroblasts showed lower rates of proliferation than did NHDFs, but ABE-treated RDEB fibroblasts showed enhanced cell proliferation compared to uncorrected cells (**Figure 2H**). Taken together, our results indicate that both ABE-mediated strategies, involving Pat1-sg1 and Pat1-sg2, are relevant for gene rescue in Pat1-derived cells.

### Prime editing for *COL7A1* gene correction in both Pat1- and Pat2-derived fibroblasts

We next investigated the potential use of prime editing for correcting the two nonsense mutations (i.e., c.2005C>T and c.3631C>T) in the *COL7A1* gene. Because PEs have a more flexible targeting scope than BEs, we sought to apply PEs for correcting both mutations. In this experiment, we used the PE3 system because of its enhanced editing efficiency compared to that of PE2. We designed pegRNAs that could correct the nonsense mutation and also induce silent mutations in the PAM sequences, because it was previously reported that such mutations in the PAM enhance the editing efficiency and reduce indel generation by inhibiting repetitive PE binding after the initial editing. We first used a pegRNA containing 13-nt PBS and 14-nt RTT together with a ngRNA. For Pat1, pegRNA#1 (Pat1-peg1, hereafter) was designed for the correction of c.3631C>T; a T-to-C conversion would occur at position +10 (10 nt downstream from the nick site) and the ngRNA would lead to the generation of a nick 60 nt downstream of the pegRNA-induced nick (**Figure 3A** and **S1B**. For Pat2, pegRNA#2 (Pat2-peg2, hereafter) was designed for the correction of c.2005C>T; in this case, a T-to-C conversion would occur at position +12 (12 nt downstream from the nick site) and the ngRNA would direct the formation of a nick in the non-edited strand at a position 56 nt upstream of the pegRNA-induced nick site (**Figure 3B** and **S1A**). Then, PE2-, pegRNA-, and ngRNA-encoding plasmids were transfected into Pat1- or Pat2-derived primary fibroblasts via electroporation. After 3 to 7 days, the cells were harvested for assessment of the editing efficiency. Genomic DNA from the bulk population of cells was subjected to high-throughput sequencing. The results showed that the average prime editing efficiencies were 10.5% at c.3631C>T with PE3/Pat1-peg1 (**Figure 3C**) and 5.2% at c.2005C>T with PE3/Pat2-peg2 (**Figure 3E**). The average indel frequencies were 1.5% for PE3/Pat1-peg1 and 0.7% for PE3/Pat2-peg2 (**Figure 3E**). When we tested various pegRNAs with different PBS lengths (i.e., 11 nt and 15 nt), pegRNAs with a 13-nt PBS led to editing activity that was comparable to that seen with the other pegRNAs (**Figure 3E**).

**Figure 3.**
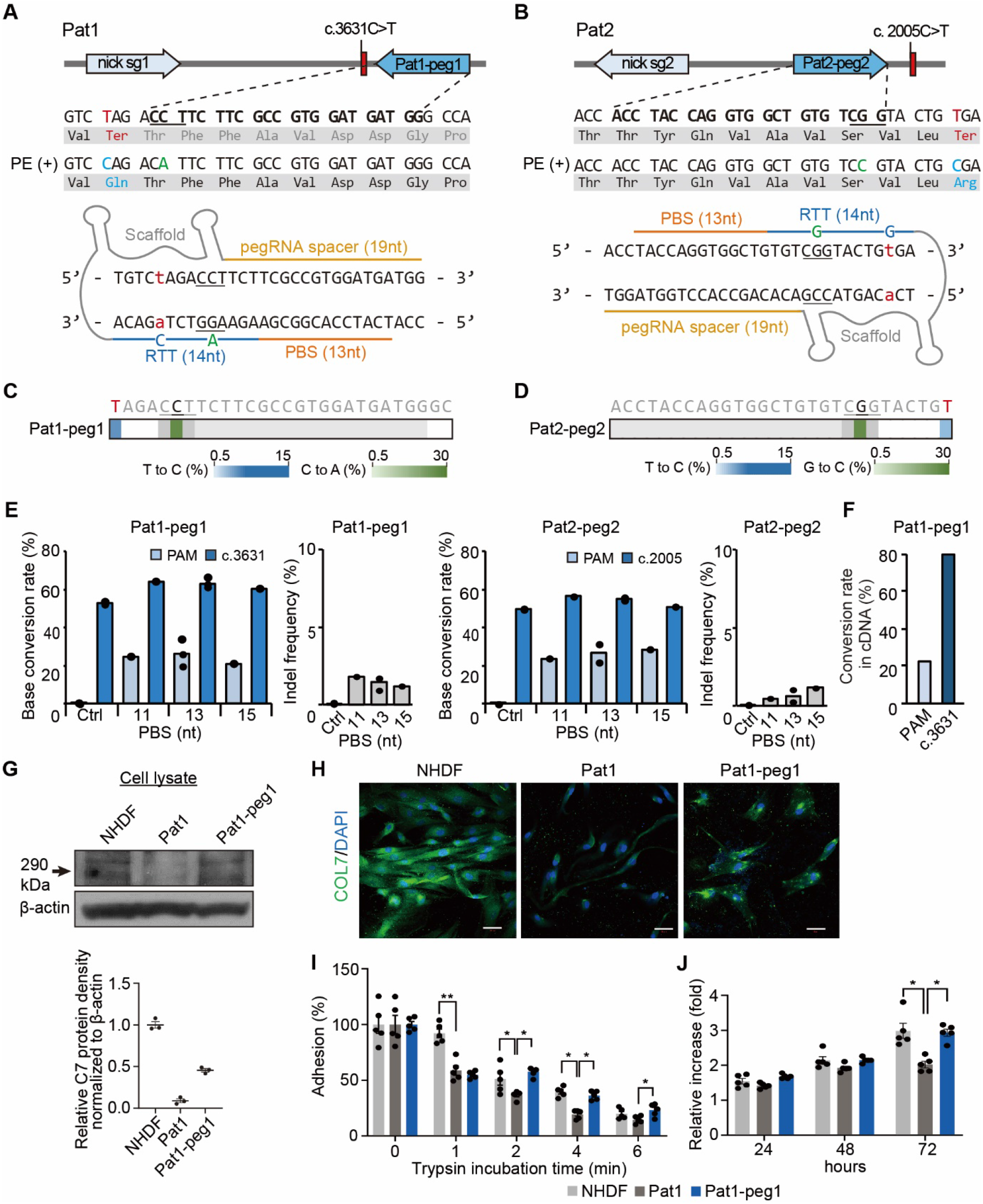
PE3-mediated correction of c.3631C>T (Q1211X) and c.2005C>T (R669X) *COL7A1* in primary fibroblasts derived from two RDEB patients. (a, b) Schematic diagram of the pegRNA and ngRNA target sites in the *COL7A1* gene in Pat1 (a) and Pat2 (b). Target sequences are highlighted in bold type, and PAM sequences are underlined. The pathogenic mutations, converted pathogenic mutations, and synonymous mutations for PAM disruption are shown in red, blue, and green, respectively (top). A 14-nt RTT and a 13-nt PBS were used for pegRNAs (bottom). (c, d) Heatmaps visualizing conversion rates determined by high-throughput sequencing. (e) Prime editing efficiencies and indel frequencies in the target sequence in the patient-derived fibroblasts transfected with various pegRNAs with different PBS lengths (n = 1∼3). (f) Conversion rates in mRNA from PE-treated RDEB fibroblasts. (g) Western blot to measure C7 abundance in cell lysates and culture supernatants using β-actin as an internal control. Protein band densities from three independent experiments are presented as bar graphs. Each density value was normalized to the β-actin value and expressed relative to the value in NHDFs. Data are mean ± SEM. (h) Immunofluorescence staining to visualize the C7 protein (green) in NHDFs, non-edited RDEB fibroblasts from Pat1, and PE3-treated RDEB fibroblasts. Nuclei were stained with DAPI (blue). Scale bars, 50 μm. (i) Trypsin-based cell detachment assay. Cell adhesion is represented as the percentage of cells that remain attached after the indicated period of trypsin treatment. Five independent experiments were performed. Data are mean ± SEM. ^*^P < 0.05, ^**^P < 0.01. (j) The proliferation of NHDFs, non-edited fibroblasts from Pat1, and PE3-treated RDEB fibroblasts was evaluated by the WST-1 assay. The ratio of the absorbance at 24, 48, and 72 hours to that at 0 hours is shown. Five independent experiments were performed. Data are mean ± SEM. ^*^P < 0.05, ^**^P < 0.01.

To investigate whether correction of *COL7A1* by the PE3 system can restore functional C7 in RDEB fibroblasts, we selected PE-treated fibroblasts derived from Pat1 because the editing efficiency was higher than that of PE-treated fibroblasts derived from Pat2. Similar to the above experiments using ABEs, we assessed the editing efficiency in cDNAs and found that the correction frequency in cDNAs was consistently higher than that in genomic DNA (**Figure 3F** and **S1C**). Western blot analysis showed that PE-treated RDEB fibroblasts expressed increased levels of the C7 protein, to up to 46% of the level in NHDFs (**Figure 3G**). In addition, immunocytochemistry confirmed efficient expression of the C7 protein in PE-treated RDEB fibroblasts, whereas the unedited cells showed no antibody reactivity (**Figure 3H**). We further evaluated the adhesion properties of the PE-treated RDEB fibroblasts by the trypsin-based cell detachment assay. PE-treated RDEB fibroblasts showed a 21%, 17%, and 7% increase in adhesion 2, 4, and 6 minutes after trypsin treatment compared to untreated RDEB fibroblasts, which showed poor cell adhesion (**Figure 3I**). We also tested the effect of *COL7A1* correction on the proliferation ability of RDEB fibroblasts using the WST-1 assay. RDEB fibroblasts carrying Q1211X and E2857X showed lower rates of proliferation than NHDFs, but PE-treated fibroblasts showed enhanced cell proliferation compared to uncorrected cells (**Figure 3J**). These findings indicate that *COL7A1* correction by PE can effectively rescue the impaired adhesion and proliferation properties of RDEB fibroblasts.

### Deposition of human type VII collagen at the DEJ of immunodeficient mice injected with ABE-/PE-corrected RDEB fibroblasts

Next, we investigated whether ABE-/PE-corrected RDEB fibroblasts could synthesize and secrete human C7 that would then localize correctly at the DEJ *in vivo* after intradermal injection into immunodeficient mice. To this end, a single dose of 5 × 10^6^ NHDFs, non-edited Pat1-derived RDEB fibroblasts, or Pat1-derived RDEB fibroblasts corrected using Pat1-sg1, Pat1-sg2, or Pat1-peg1 suspended in 150 μl of phosphate-buffered saline was intradermally injected into the back skin of immunodeficient mice (**Figure 4A**). Two weeks after injection, human C7 protein deposition at the DEJ was analyzed by immunofluorescence using anti-human COL7 antibody (kindly provided by Dr Hiroaki Iwata, Hokkaido University Graduate School of Medicine). We found that skin injected with ABE-/PE-treated RDEB fibroblasts showed strong linear deposition of human C7 along the DEJ, whereas human C7 was barely detectable in skin injected with phosphate-buffered saline alone or uncorrected RDEB fibroblasts (**Figure 4B**). These observations clearly demonstrate that ABE- or PE-mediated correction of a *COL7A1* nonsense mutation functionally restores the expression and secretion of C7 in primary RDEB fibroblasts.

**Figure 4.**
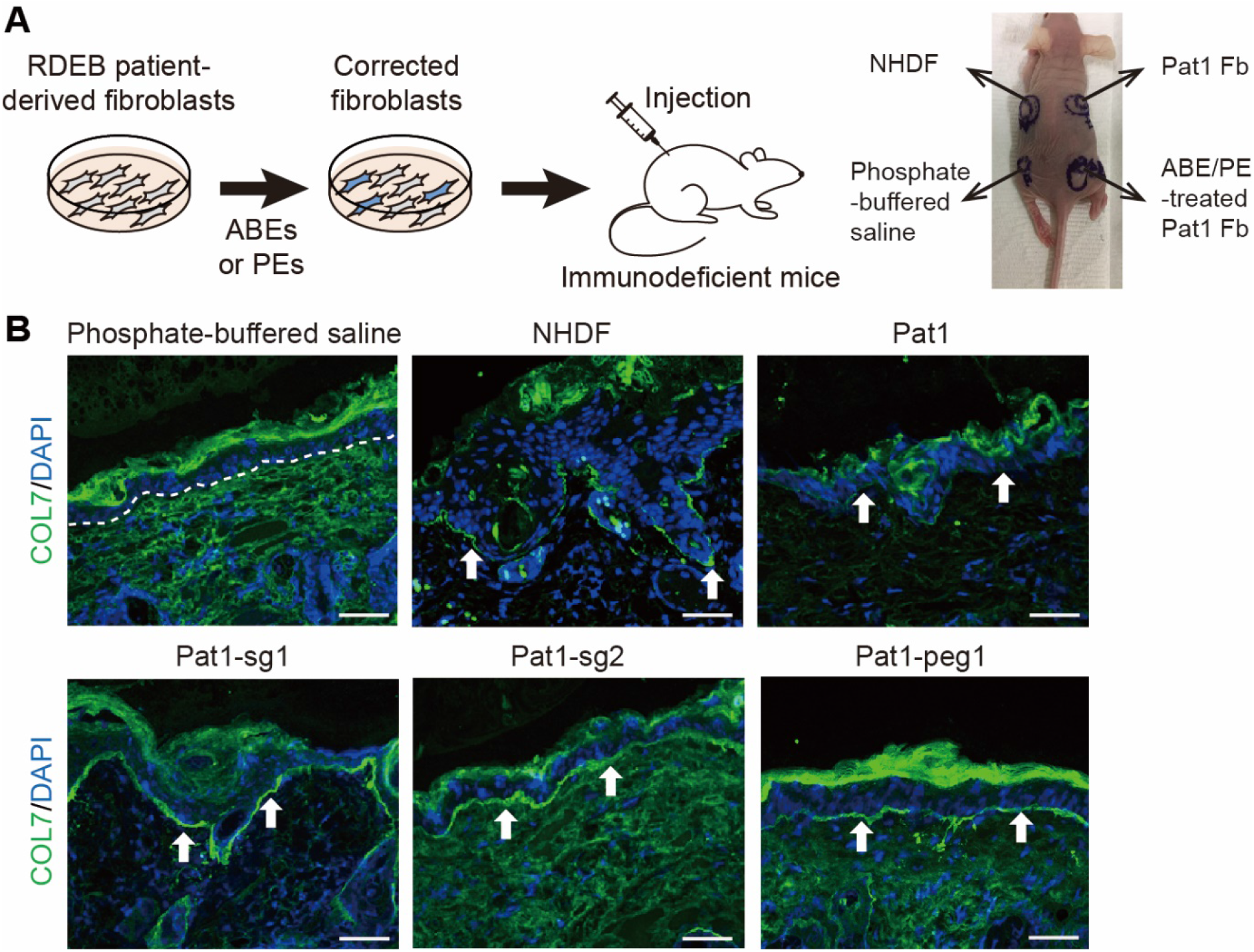
Correct deposition of human C7 at the DEJ in immunodeficient mice after intradermal injection of ABE- or PE-treated primary RDEB fibroblasts. (a) Scheme of the experiment in which ABE-/PE-treated patient-derived fibroblasts were injected into the mouse model. (b) Immunofluorescence staining to visualize the C7 protein. NHDFs, non-edited fibroblasts from Pat1, ABE-treated RDEB fibroblasts, PE-treated RDEB fibroblasts (5×10^6^ cells/150 μl of phosphate-buffered saline), or phosphate-buffered saline were intradermally injected into the back skin of immunodeficient mice. Two weeks after the injections, immunofluorescent analysis of C7 (green) was performed using a rabbit polyclonal antibody that recognizes human C7. The white dotted line indicates the DEJ. White arrows indicate human C7 deposited at the DEJ. Scale bars, 50 μm. Fb, fibroblasts; NHDFs, normal human dermal fibroblasts.

We also investigated whether off-target DNA editing occurred in Pat1-derived RDEB fibroblasts corrected using Pat1-sg1, Pat1-sg2, or Pat1-peg1. We carefully identified potential off-target sites using Cas-OFFinder software.^[24]^ When up to three mismatched bases or one mismatched base with a DNA/RNA bulge were allowed, a total of six potential off-target sites for Pat1-sg1, no sites for Pat-sg2, and seven sites for Pat1-peg1 were identified. High-throughput sequencing revealed that no off-target editing was found in any of the cell populations (**Figure S2**).

## DISCUSSION

Genome editing has emerged as a promising molecular approach for treating genetic diseases. In this study, we first established a *COL7A1* mutation database containing 69 pathogenic mutations (40 variants without recurrence) from a total of 36 Korean RDEB patients. As a proof-of-concept, we chose two patients having two representative *COL7A1* mutations [i.e., c.3631C>T (Q1211X) and c.2005C>T (R669X)], which were formerly shown to cause generalized severe RDEB^[18b, 19]^ and have been reported to result in nonsense-mediated mRNA decay that manifests as a complete absence of C7.^[18]^ We first applied ABEs to correct the mutations in primary RDEB patient-derived fibroblasts through two different strategies: direct correction of target mutations to the wild-type sequence and induction of readthrough of a PTC with the CRISPR-pass method. After electroporation of the plasmid-based delivery system, ABEmax showed an editing efficacy (24.6∼30.6% in genomic DNA and 37∼58% at the mRNA level) comparable to that in previous work by Osborn et al. in which the ABEmax-induced gene correction rate was 8.2%∼23.8% in genomic DNA and 17.8∼45% at the mRNA level in primary RDEB fibroblasts.^[15]^ In addition, ABEmax induced a low rate of indels (0.1∼3.3%), similar to that in the previous research (1.5∼1.9%),^[15]^ suggesting a more reliable editing approach than HDR-mediated gene correction, which resulted in higher indel rates. This point is important because of DSB-associated safety issues.

However, BEs cannot be used to correct the disease-associated mutations in more than half of RDEB patients. Furthermore, we observed that the frequency of ABE-induced bystander edits varied over a wide range, with an upper value of 44.5% (1.8-44.5%), depending on the width of the editing window. This effect might limit further applications of BEs for RDEB treatment. As a potential alternative, we next used PEs to correct the mutations. To the best of our knowledge, this study is the first to demonstrate the feasibility of PEs for correcting pathogenic *COL7A1* mutations, including a mutation that was not suitable for correction with an ABE recognizing the canonical SpCas9 PAM, to treat RDEB. Although the editing efficiencies of PE3 on the compound heterozygous Q1211X (10.5%) and R669X (5.2%) targets in primary RDEB patient-derived fibroblasts were overall lower than that of ABEs, PEs showed precise base correction with few bystander edits. In addition, PE3 did not induce detectable off-target editing at potential Cas9 off-target loci, consistent with previous observations.^[16, 25]^

We further found that correction of these two nonsense mutations by either ABE or PE restored the synthesis and secretion of full-length C7 in RDEB fibroblasts. ABE- and PE-mediated genetic correction also rescued the poor adhesion capacity and growth potential of RDEB fibroblasts. We ultimately observed that ABE- and PE-treated RDEB fibroblasts produced functional C7 that was correctly deposited into the DEJ at the site of injection into the skin of immunodeficient mice. It has been shown that C7 levels that are 35% of that in NHDFs are sufficient to provide mechanical stability of the skin in a DEB hypomorphic murine model.^[26]^ In this study, ABE- and PE-mediated correction of Q1211X in RDEB fibroblasts respectively restored the production of full-length C7 to levels of up to 68% and 46% of that in NHDFs; furthermore, the C7 was correctly deposited along the DEJ of the immunodeficient mouse skin. Edited RDEB fibroblasts showed enhanced proliferation compared to non-edited cells, which may explain why the levels of C7 restoration are higher than would be expected from the editing frequency at the genomic DNA level.

Collectively, our data demonstrate that both ABEs and PEs enable efficient correction of pathogenic *COL7A1* mutations with higher ratios of the desired edit per indel than HDR in patient-derived primary cells, restore C7 expression to levels known to rescue the phenotype of the DEB murine model, and induce the formation of functional C7 that incorporates into the DEJ of immunodeficient mice. By using a non-viral delivery method, electroporation, we minimize safety concerns for therapeutic translation of these editing technologies to treat RDEB. Despite the higher editing efficacy of ABEs compared to PEs for correcting *COL7A1* nonsense mutations, PEs would be more reliable tools than ABEs for RDEB treatment, considering that PEs exhibit precise correction without bystander edits, the ability to target almost all pathogenic mutations, and negligible off-target editing effects. In the near future, we expect that BEs or PEs will be used for treating RDEB patients *via* the transplantation of *ex vivo* gene-corrected autologous cells.

## ACKNOWLEDGEMENTS

This research was supported by grants from the National Research Foundation of Korea (NRF) no. 2021R1A2C3012908 and no. 2021M3A9H3015389 to S.B, and no. 2021R1A2C201370 to S.E.L.

## AUTHOR CONTRIBUTIONS

S.B. and S.E.L. conceived this project; S.-E.K., S.-A.H., and A.-y.L. performed and analyzed the experiments; G.-H.H. performed bioinformatics analyses; J.H.K., H.I., and S.-C.K. gave critical comments; S.-A.H., S.B., and S.E.L. wrote the manuscript with the approval of all other authors.

## Additional information

Supplementary Information accompanying this paper is available at http://

## Methods

### Analysis of targetable disease mutations in *COL7A1*

RDEB-associated variants were collected from the *COL7A1* variants database (http://www.col7a1-database.info). The information of reference sequence and CDS position of each variant were obtained from the National Center for Biotechnology Information (NCBI) website (NG_007065). The number of possible BE-targetable variants was calculated when the mutations were located within editing activity windows; 3^rd^ to 9^th^ positions counting from the 5’ end of the target sequence. The number of possible PE-mediated gene corrections was counted when distances between the mutations and Cas9-mediated nick sites were 12 bp or less. The analysis program was developed using using Python3.

### Establishment of a *COL7A1* mutation database specific for Korean RDEB patients

RDEB patients from all regions of South Korea were referred to Gangnam Severance Hospital, Seoul, Korea, for molecular diagnosis. The results, which include information about 38 patients from 35 unrelated families, make up the largest Korean database for RDEB. RDEB was diagnosed based on clinical features, immunofluorescence antigen mapping and next-generation sequencing (NGS) and/or Sanger sequencing of *COL7A1*. All participants or their legal guardians gave their written informed consent, and this study was approved by the Institutional Review Board (IRB) at Gangnam Severance Hospital in accordance with the principles of the Declaration of Helsinki. Genomic DNA was extracted from peripheral blood lymphocytes from patients. DNA from five families was analyzed by NGS, and DNA from 30 families was analyzed by traditional Sanger sequencing. All 118 *COL7A1* exons and exon– intron borders were amplified by polymerase chain reaction (PCR) and the products were subsequently sequenced. For all mutations other than nonsense mutations, 100 control alleles were studied to rule out the possibility that the putative disease-associated mutation might be a frequent polymorphism.

### Study approval and human subjects

Two patients with RDEB carrying compound heterozygous *COL7A1* mutations (c.3631C>T, p.Q1211X, exon 27 and c.8569G>T, p.E2857X, exon 116 in patient 1; c.2005C>T, p.R669X, exon 15 and c.8569G>T, p.E2857X, exon 116 in patient 2) were enrolled in this study approved by the Gangnam Severance Hospital IRB (no. 3-2021-0485). Declaration of Helsinki protocols were followed, and both subjects gave written informed consent for the donation of skin cells.

### Isolation and culture of primary cells from patients with RDEB

Skin samples from the RDEB patients, which were obtained by 3-mm punch biopsies, were dissected into 10 pieces with sharp scalpels. For skin explant culture, the pieces were placed in and attached to the well of a 100-mm dish and maintained at 37°C with 5% CO_2_ in Dulbecco’s modified Eagle medium (DMEM), supplemented with 10% fetal bovine serum (FBS) and 1% penicillin/streptomycin (P/S). After 1 week, the medium was changed to Keratinocyte-SFM medium (Thermo Fisher Scientific) supplemented with 1% P/S for keratinocyte culture and DMEM supplemented with 10% FBS and 1% P/S for fibroblasts. Primary human keratinocytes and fibroblasts were cryopreserved at the second passage and stored at -80°C until use.

### Sanger sequencing

Genomic DNA was extracted from patient blood samples using ExgeneTM Blood SV mini (GeneAll, Seoul, South Korea). The *COL7A1* gene was amplified by PCR using targeted primers (**Table S2**).

### Immunofluorescence for C7 in human skin\

Frozen skin tissues from a normal individual and the patients were sectioned at 6 μm and stained with mouse monoclonal anti-NC1 C7 antibody (clone LH 7.2; Sigma-Aldrich) at a 1:1000 dilution. Alexa Fluor 488 conjugated rabbit anti mouse IgG (Thermo Fisher Scientific) was used as secondary antibody. Sections were stained with 4,6-diamidino-2-phenylindole (DAPI) (Thermo Fisher Scientific). Images were captured using an LSM 780 confocal microscope (Carl Zeiss, Oberkochen, Germany).

### Western blots

Total proteins from primary fibroblasts were isolated using RIPA buffer (Cell Signaling Technology, Danvers, MA) supplemented with 1 mM phenylmethylsulfonyl fluoride (PMSF). The fibroblast culture supernatants were mixed with acetone and centrifuged at 4000 rpm for 20 minutes, after which the resulting pellets were washed with phosphate-buffered saline. Total proteins from these supernatant-derived pellets were isolated using RIPA buffer (Cell Signaling Technology, Danvers, MA) supplemented with 1 mM PMSF. After protein isolation, equal amounts of proteins from each group were loaded onto Nupage Novex Bis-Tris Gels (Thermo Fisher Scientific), and electrophoresis was performed using an X-cell SureLock Mini-Cell (Thermo Fisher Scientific). After electrophoresis, proteins were transferred onto polyvinylidene difluoride membranes, which were then incubated with rabbit polyclonal anti-collagen VII antibody (ab93350; Abcam) that was diluted in Tris-buffered saline (TBS) containing 0.05% Tween 20 (TBS-T), at a dilution of 1:1000. Blots were washed with 0.05% TBS-T and then incubated with horseradish peroxidase-conjugated anti-mouse and anti-rabbit secondary antibodies (Thermo Fisher Scientific) in 0.05% TBS-T at a dilution of 1:2000. Blots were developed using ECL PLUS reagent (Pierce, Rockford, IL). The densities of the resulting protein bands were analyzed using ImageJ densitometry software (National Institutes of Health, Bethesda, MD).

### Immunocytochemistry

For immunocytochemistry, fibroblasts were cultured in chamber slides (LabTek, Thermo Fisher Scientific), fixed with 4% paraformaldehyde for 10 minutes, blocked with 0.5% bovine serum albumin for 30 minutes, and then incubated with rabbit polyclonal anti-collagen VII antibody (ab93350; Abcam; 1000-fold dilution) overnight at 4°C. After washing, cells were incubated with Alexa Fluor 488-conjugated rabbit anti-mouse IgG (Thermo Fisher Scientific) as the secondary antibody at a 1:2000 dilution and DAPI (Thermo Fisher Scientific). Images were captured using an LSM 780 confocal microscope (Carl Zeiss, Oberkochen, Germany).

### Intradermal injection of RDEB fibroblasts into immunodeficient mice

All animal experiments were approved by the Animal Care Committee of Yonsei University College of Medicine. Male athymic nude mice (nu/nu) (Central Lab Animal Inc., Seoul, Korea) were maintained under specific pathogen-free conditions with water, food, and supportive nutrition ad libitum. Three fibroblast populations (NHDFs, RDEB fibroblasts, and ABE- and PE-treated RDEB fibroblasts) were expanded to obtain the required number of cells for intradermal injections. Then, cells were harvested using Trypsin/ethylenediaminetetraacetic acid (EDTA) (Life Technologies), after which they were washed gently three times with phosphate-buffered saline. Five million of each fibroblast type were resuspended in 150 μl of phosphate-buffered saline and were intradermally injected with a 24 G needle in a 1 cm^2^ area. A single 150 μl volume was delivered via two injections of 75 μl. Three mice were injected per group. Two weeks after injection, mouse skin samples were obtained for immunofluorescence staining for human C7.

### Immunofluorescence staining of mouse skin

For immunofluorescence detection of human C7 in injected mouse skin, frozen skin tissues were sectioned at 6 μm and stained with polyclonal rabbit anti-COL7 antibody (anti-FNIII7-FNIII8 antibody, kindly provided by Dr. Hiroaki Iwata, Department of Dermatology, Hokkaido University Graduate School of Medicine, Sapporo, Japan), at a dilution of 1:1000 at 4°C overnight. Alexa Fluor 488-conjugated rabbit anti-mouse IgG (Thermo Fisher Scientific) was used as the secondary antibody. Sections were stained with DAPI (Thermo Fisher Scientific). Images were captured using an LSM 780 confocal microscope (Carl Zeiss, Oberkochen, Germany).

### Cell detachment assay

Fibroblasts were seeded at a density of 6 × 10^4^ cells/well in a 96-well plate and cultured for 24 hours. After washing with 1X phosphate-buffered saline, confluent layers of fibroblasts were treated with 0.05% trypsin/EDTA for 6, 4, 2, 1, and 0 minutes, followed by washing once with 10% FBS/DMEM to inactivate trypsin and then twice with phosphate-buffered saline. The adherent cells were stained with 0.5% crystal-violet (Sigma Aldrich) for 30 minutes and lysed with 1% sodium dodecyl sulfate (Sigma Aldrich). The percentage of adherent cells was determined by measuring the absorbance at 590 nm using a spectrophotometer.

### Proliferation assay

NHDFs, unedited RDEB fibroblasts, and corrected RDEB fibroblasts were seeded at a concentration of 5 × 10^3^ cells/well into microplates (tissue culture grade, 96 wells, flat bottom) in 100 μl 10% FBS/DMEM culture medium per well. 24 hours, 48 hours, or 72 hours after incubation at 37°C with 5% CO_2_, cellular proliferation was evaluated using a WST-1 assay (05015944001, Roche, Basel, Switzerland). Briefly, the cells were incubated with the WST-1 reagent for 4 hours, and absorbance at 450 nm and 650 nm (reference wavelength) was detected using a microplate reader (iMark, Bio-Rad).

### Construction of sgRNA- and pegRNA-expressing plasmids

The target sequences were selected using Cas-designer (http://www.rgenome.net/cas-designer/).^[27]^ To construct sgRNA- and ngRNA-expressing plasmids, complementary oligos representing target sequences were annealed and cloned into pRG2 (Addgene #104174). To construct pegRNA-expressing plasmids, complementary oligos representing target sequences, the sgRNA scaffold, and 3’ extensions were annealed and cloned into pU6-pegRNA-GG-acceptor (Addgene #132777). The oligos are listed in **Table S3**.

### Transfection

Electroporation was performed using a Neon Transfection System (Thermo Fisher) with the following parameters: voltage, 1,650; width, 10ms; number, 3. For base editing, 150,000 patient-derived fibroblasts were transfected with 900 ng of ABEmax-encoding plasmid (Addgene, #112095). For prime editing, 150,000 patient-derived fibroblasts were transfected with 900 ng of PE2-encoding plasmid (Addgene #132775), 300 ng of pegRNA-encoding plasmid, and 83 ng of ngRNA-encoding plasmid.

### Cell lysis and high-throughput sequencing

Cell pellets were resuspended in Proteinase K extraction buffer [40 mM Tris-HCl (pH 8.0) (Sigma), 1% Tween 20 (Sigma), 0.2 mM EDTA (Sigma), 10 mg of Proteinase K, 0.2% nonidet P-40 (VWR Life Science)] and then incubated at 60°C for 15 minutes and 98°C for 5 minutes. 1∼3 μL of Proteinase K extraction solution containing genomic DNA was amplified for high-throughput sequencing. ABE and PE target sites were amplified using SUN-PCR blend (Sun Genetics). The PCR products were purified using Expin™ PCR SV mini (GeneAll) and sequenced using a MiniSeq Sequencing System (Illumina). The results were analyzed using Cas-Analyzer (http://www.rgenome.net/cas-analyzer/), BE-Analyzer (http://www.rgenome.net/be-analyzer/), and PE-analyzer (http://www.rgenome.net/pe-analyzer/).^[28]^ The primers are listed in **Table S2**.

## Data Availability

High-throughput sequencing data have been deposited in the NCBI Sequence Read Archive database (SRA; https://www.ncbi.nlm.nih.gov/sra) under accession number PRJNA739484.

## Code availability

The authors declare that all unreported custom Python code used in this study is available from the corresponding author upon reasonable request.

